# Prolonged ovarian storage of mature *Drosophila* oocytes dramatically increases meiotic spindle instability

**DOI:** 10.1101/675389

**Authors:** Ethan J. Greenblatt, Rebecca Obniski, Claire Mical, Allan C. Spradling

**Author notes:** ^*^Author for correspondence, Corresponding Author: Dr. Allan C. Spradling.

## Abstract

More than 95% of fertilized *Drosophila* oocytes from outbred stocks develop fully regardless of maternal age, in contrast to human oocytes, which frequently generate non-viable aneuploid embryos. Since *Drosophila* oocytes are normally stored only briefly prior to ovulation, unlike their human counterparts, we investigated the effects of storage on oocyte quality. Using a novel system to acquire oocytes held for known periods, we analyzed by ribosome profiling how translation and cellular function change over time. Oocyte developmental capacity decays in a precise temperature-dependent manner over 1-4 weeks, due to a progressive inability to complete meiosis. Meiotic metaphase genes, the *Fmr1* translational regulator, and the small heat shock protein chaperones *Hsp26* and *Hsp27* are preferentially translated during storage, and oocytes lacking *Hsp26* and *Hsp27* age prematurely. However translation falls generally 2.3-fold with age despite constant mRNA levels, and this inability to maintain translational equilibrium correlates with oocyte functional decline. These findings show that meiotic chromosome segregation in *Drosophila* oocytes is uniquely sensitive to prolonged quiescence, and suggest that the extended storage of mature human oocytes contributes to their chromosome instability. If so, then these problems may be more amenable to intervention than previously supposed.

## Introduction

Animal oocytes grow extensively to become the largest non-polyploid body cells, but at a few specific stages ovarian follicles can persist in a non-growing state. Following recombination and prophase arrest at the diplotene stage of meiosis, mammalian oocytes within primordial follicles cease demonstrable development to establish the “ovarian reserve,” whose slow utilization over multiple decades in humans determines the duration of female fertility. Both oocytes and granulosa cells within primordial follicles are bathed in maternal nutrients and remain able to transcribe and translate genes. If these non-growing cells use any special strategies to achieve extraordinary longevity they remain poorly known. Eventually, arrested oocytes resume growth and develop to their final size while remaining in meiotic diplotene. Shortly before fertilization, oocytes mature, during which meiosis resumes and progresses to an arrest at metaphase I or II (Coticchio et al., 2015; Hughes et al., 2018).

In many species, fully grown oocytes also have a period of quiescence. In a good nutritional environment, mated *Drosophila* females ovulate mature oocytes shortly after they reach their final size. However, *Drosophila* store metaphase I-arrested oocytes for multiple days if food or sperm are unavailable, despite a lack of transcription. Analysis of polysomes and translational activity also suggested that stored oocytes maintain protein production, though at a reduced level (Kronja et al., 2014a; Lovett and Goldstein, 1977). Likewise, mammalian oocytes routinely cease transcription upon reaching full size (ABE et al., 2010; Jukam et al., 2017). Oocytes remain transcriptionally inactive until zygotic genome activitation at the two-cell stage (mouse) or at the 4-cell stage (human). Previously it has been difficult to study the functional and molecular implications of mature oocyte storage because obtaining pure populations of oocytes maintained as primordial follicles or as mature oocytes for known periods of time was not possible.

Storing oocytes is generally associated with a significant risk of functional impairment in mammals. In humans, where all oocytes are stored to some extent, a portion of oocytes develop meiotic segregation errors that are the major cause miscarriage. Past the age of 35, chromosome mis-segregation further increases as reflected in exponentially growing rates of Down’s syndrome (Webster and Schuh, 2017). The high frequency of meiotic defects in human oocytes has been explained by the exceptional length of time they spend as arrested primordial follicles (Chiang et al., 2010; Herbert et al., 2015). However, it has not been ruled out that chromosome segregation defects may also result from storage after completion. Mature human oocytes can only be maintained in culture for a brief period (Alikani et al., 2000). In addition, studies of *in vitro* fertilized human oocytes suggest that spindle-related errors in mitotic chromosome segregation during early embryonic cell cycles are frequent even in embryos derived from donor eggs of young women (McCoy et al., 2015).

Here we show that mature *Drosophila* oocytes remain capable of supporting embryonic development for many days while stored in the ovary, providing a system for the molecular genetic analysis of oocyte aging. Oocytes stored only briefly develop with high fidelity. However, as aging continues, completing meiosis successfully following fertilization becomes the major factor limiting oocyte viability. Cytologically detectable spindle defects increase during storage and early developmental arrest gradually become the predominant fate of the resulting embryos. Translation of mRNAs encoding meiotic metaphase and spindle-related proteins decline as part of a general 2.3-fold reduction during aging in the absence of bulk changes to mRNA levels. Our findings show that storage of highly functional mature oocytes *in vivo* is sufficient to destabilize chromosome segregation, suggesting that the prolonged storage of mature oocytes may be an important source of meiotic chromosome instability in human females.

## Results

### A general method for studying *Drosophila* oocyte aging

*Drosophila* ovaries are organized into highly regulated ovarioles that preserve the order in which follicles develop (Fig. 1A). Ovarian biology allowed us to develop a method to obtain mature oocytes that have been stored in the ovary for a known period of time. Newly eclosed virgin female flies with immature ovaries are fed a nutrient-rich yeast paste that stimulates exactly two young follicles per ovariole to develop past a nutrient-sensitive checkpoint at stage 8 (Fig. 1A,B). Withdrawal of the yeast food after 24 hours prevents any additional follicles from passing the checkpoint. However, the two oldest follicles grow to maturity (stage 14). In the absence of mating, these mature eggs are stored in the ovary indefinitely and not replaced, as shown by the continuing absence of post-checkpoint stage 10 oocytes (Fig. 1B; Greenblatt and Spradling, 2018). The stored mature oocytes will be ovulated first when egg laying is finally stimulated, whereas developing oocytes will require at least 24 hours after stimulation to finish developing from stage 8 to stage 14.

**Figure 1.**
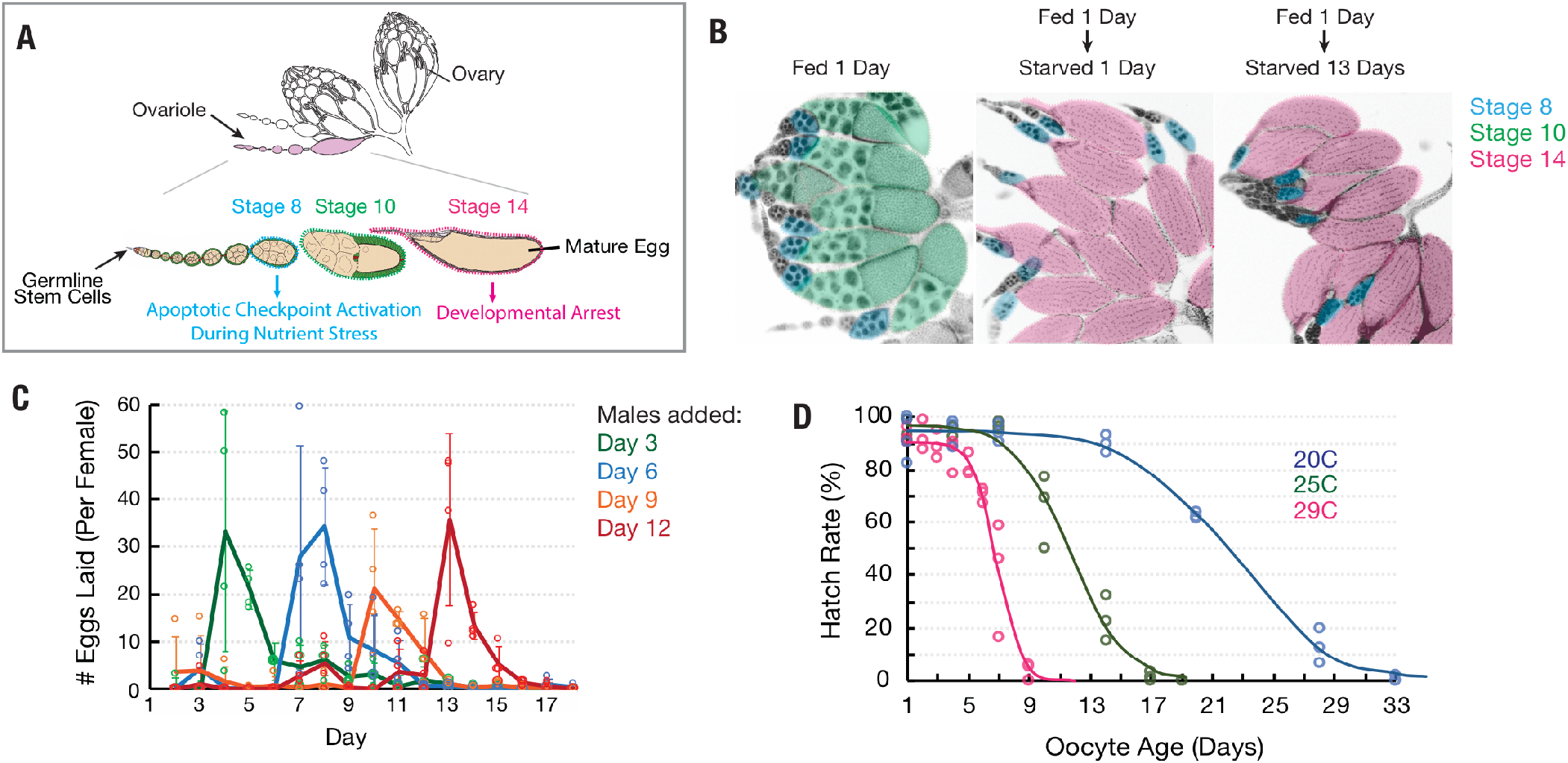
Oocytes age reproducibly in a temperature-dependent manner. (A) A schematic of the ovarioles that make up a Drosophila ovary (above) and the structure of a single ovariole (below) showing the germarium (left) followed by seven increasingly mature follicles. Stages 8, 10 and a terminal, mature stage 14 follicle are labeled. (B) DAPI stained ovaries from females that were fed 1 day (left), fed 1 day then deprived of yeast (“starved”) 1 day (middle), and fed 1 day then starved 13 days (right). Oocytes of the three stages shown in A are colored according to developmental stage, revealing the stable storage of two mature stage 14 oocytes per ovariole. (C) Egg laying data for females subjected to the storage protocol, and then provided with males after 3 (green), 6 (blue), 9 (orange) or 12 (red) days. In all four cases, mating stimulates the release of the previously stored stage 14 oocytes as fertilized embryos. (D) Aging curves (days) for follicles stored *in vivo* at 29°C (magenta), 25°C (green) or 20°C (blue). For each point, virgin females retaining oocytes for the indicated number of days at the indicated constant temperature were crossed to males and at least 100 laid embryos were followed and the percentage undergoing normal development to hatching determined.

We measured the stability of stored oocytes over time by placing females with held, mature oocytes of known age with males, which stimulates the rapid fertilization and deposition of their mature oocytes following mating (Fig. 1C). We found that while young oocytes support development to hatching at high rates (90-96%), older oocytes generate embryos with lower levels of hatching. Loss of developmental capacity follows highly reproducible sigmoidal kinetics over the course of 1-4 weeks depending on temperature (Fig. 1D). At 29°C, 25°C or 20°C, 50% of eggs fail to hatch after about 7, 12, or 23 days of storage, respectively (Fig. 1D). Thus, by appropriately feeding newly eclosed females and delaying mating, we are able to obtain oocytes for study of known age that are at a known point on an aging curve.

### Protein translation declines during oocyte aging

In order to investigate the gene products actively translated by mature oocytes as they aged, we isolated 2, 8, and 12-day old stage 14 follicles (at 25°C), whose hatch rates are 96%, 89%, and 44%, respectively, removed their follicle cells (see Methods), and performed mRNA-seq and ribosome profiling in triplicate. We added a constant amount of ovarian lysate from *D. pseudoobscura (Dpse)* as a spike-in to allow us to measure changes in translation globally as well as at the single gene level. *Dpse* is sufficiently diverged from *D. melanogaster (Dmel)* that >97% of 30 nucleotide ribosome footprints contain at least one polymorphism that can be distinguished by sequencing (see Methods). Ribosome profiling experiments were highly reproducible (Fig. S1A-D,F). Total mRNA levels, as determined by the ratio of *Dmel* to *Dpse* sequencing reads, did not change significantly between 2 and 12 days of oocyte age (Fig. 2A). Not only the amount of mRNA, but mRNA composition was also unchanged during aging. RNA-seq transcripts per million (TPM) values from oocytes at day 2 or day 12 of aging correlated very strongly (Fig. 2B). By contrast, bulk translation in 8 and 12 day old oocytes was reduced to 59% and 43%, respectively, of levels in 2 day oocytes (Fig. 2C). Thus, the overall level of translation declines significantly as oocytes age in vivo over a ten-day period.

**Figure 2.**
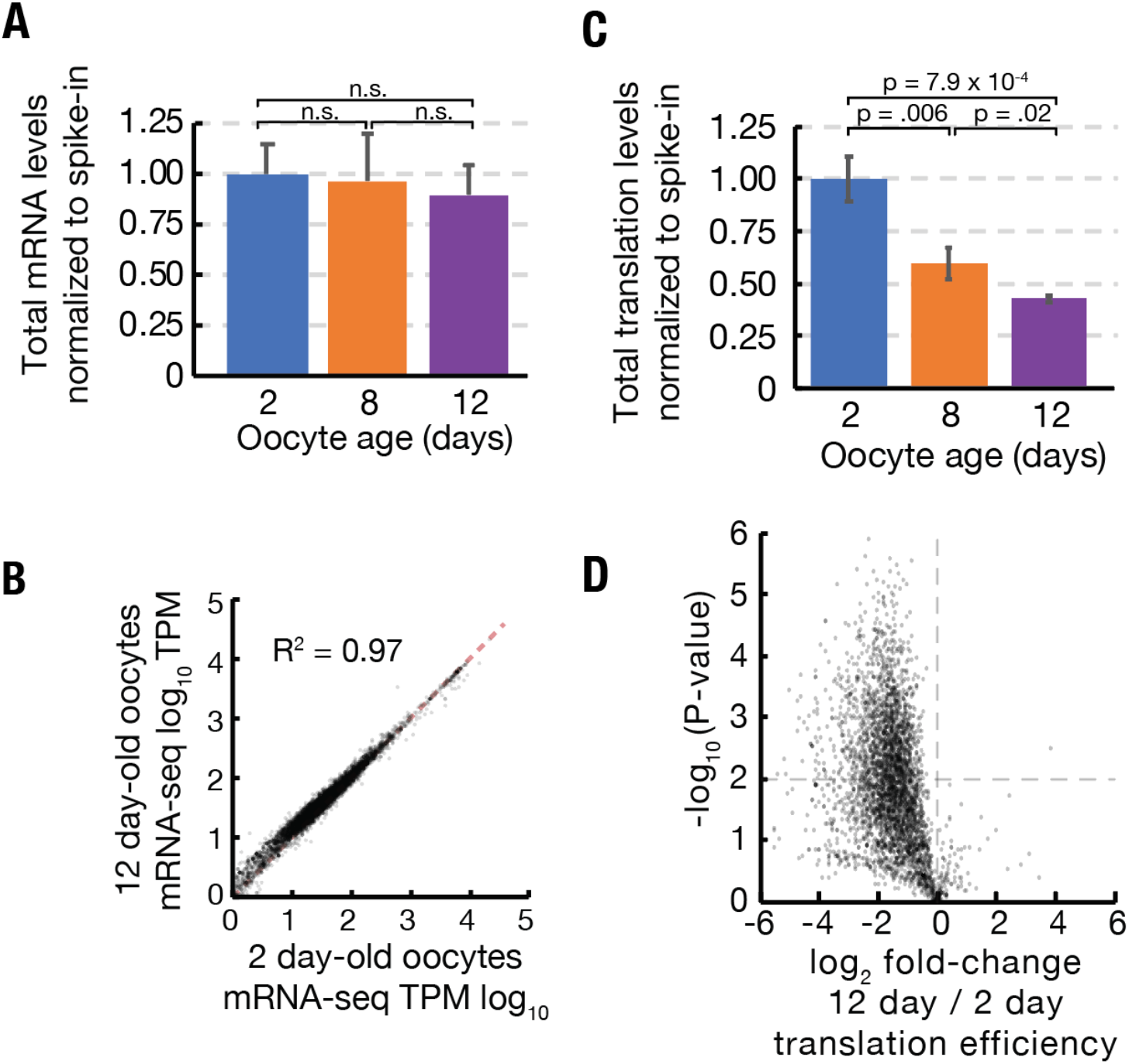
Stability of mRNA levels and translation of stored stage 14 follicles. (A) Average mRNA levels per mature follicle normalized to spike-in control in stage 14 oocytes from females aged for 2 (blue), 8 (orange) or 12 (purple) days. Differences are not significant (Student’s t-test). (B) Total translation levels per mature follicle normalized to spike-in control were compared between oocytes aged for 2 (blue), 8 (orange) or 12 (purple) days. Differences are significant as shown (Students t-test). (C) Log-Log plot showing high correlation (R^2^ = 0.97) of mRNA-seq values (TPM, transcripts per million) from stage 14 follicles retained at 25°C for 2 days (96% viability) vs 12 days (44% viability). Equal expression (dashed red line). (D) Volcano plot showing global reduction in translation efficiency in 12 day versus 2 day oocytes.

### Aging reduces the translation of all classes of genes

In order to determine whether declining translation in aging oocytes was a general phenomenon or preferentially affected a subset of genes, we analyzed changes to translation and mRNA levels from individual genes. The translation of germline expressed genes such as *Hsp26*, *vtd*, and *me31B*, but not their mRNA levels, reproducibly declined, much like total translation (Fig. 3A). The changes in translation were not caused by premature egg activation. Genes such as *CycB*, *CycA*, *and bora*, whose translation is substantially upregulated at the start of embryogenesis did not increase in aged oocytes (Fig. 3B). Rather, translational efficiency declined globally (Fig. 2D).

**Figure 3.**
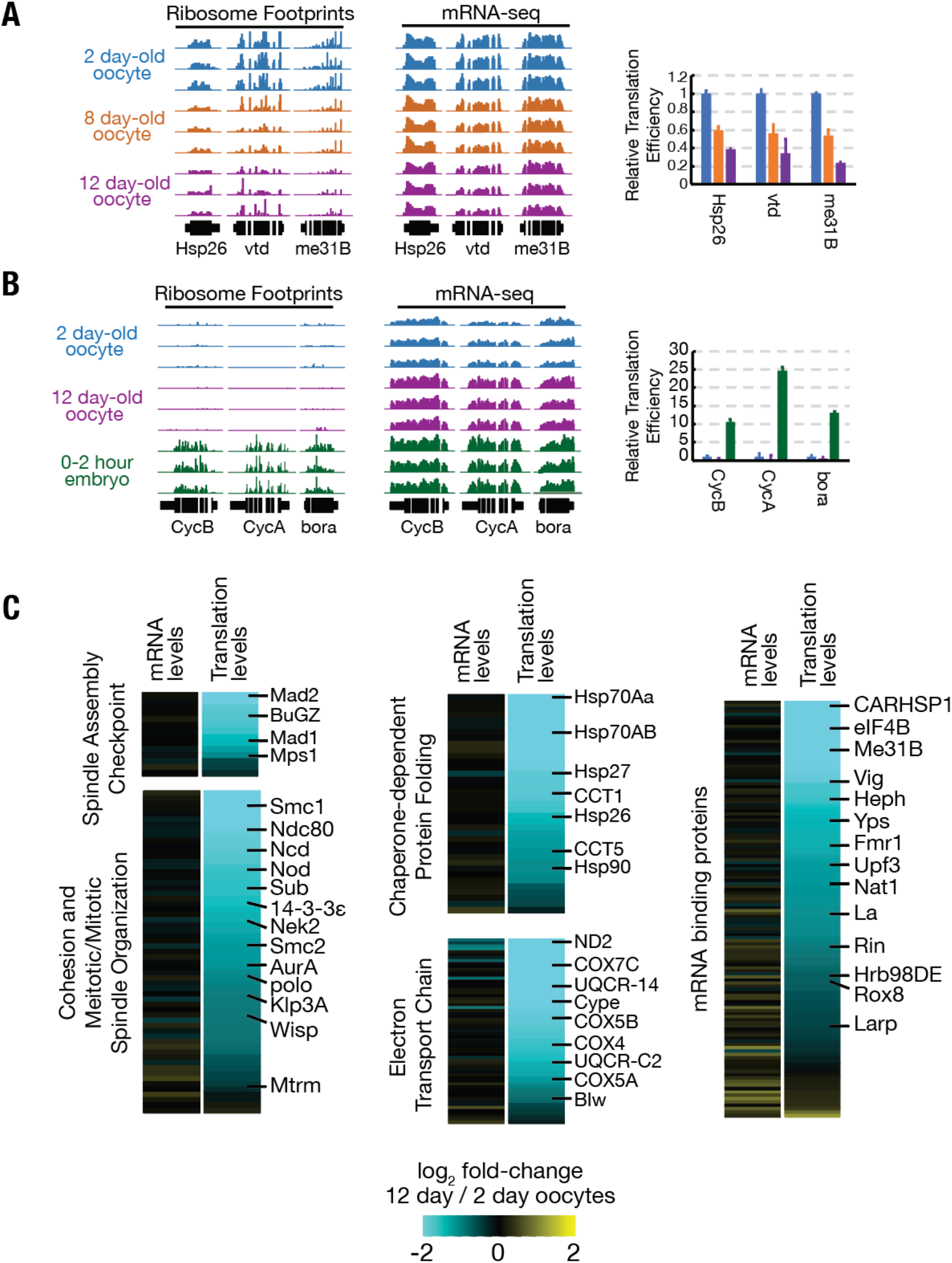
Translation is reduced globally during oocyte aging. (A) Relative read depths from replicate ribosome footprinting and mRNA-sequencing experiments of *Hsp26*, *vtd*, and *me31B* from 2, 8, and 12 day old oocytes. Data were normalized to spike-in controls. Relative translational efficiency (right panel) falls 2.5-5 fold between day 2 and day 12. (B) Read depths and translational efficiency values are shown as in (A) for genes preferentially translated in embryos *CycB*, *CycA*, and *bora* from 2 day oocytes, 12 day oocytes, and 0-2 hour embryos from non-aged oocytes. (C) Heat maps showing reduced translation, but similar mRNA levels, of genes of various GO categories in 12 day oocytes as compared to 2 day oocytes. Specific genes are indicated.

We grouped genes in several functional categories with potential relevance to oocyte aging and compared changes in the mRNA levels and translation levels (Fig. 3C). Widespread reductions were seen among genes with the GO categories spindle assembly checkpoint, meiotic/mitotic spindle organization, chaperone-dependent protein folding, electron transport chain, and mRNA binding proteins (Fig. 3C). For each category, the location of several well-known genes is indicated (Fig. 3C), including genes shown previously to be dose sensitive for chromosome stability, such as *sub*, *ncd*, *nod*, and *SMC1* (Knowles and Hawley, 1991; Moore et al., 1994; Subramanian and Bickel, 2008; Zhang et al., 1990).

### A small subset of mRNAs are translationally upregulated in arrested mature oocytes

Mature oocytes store mRNAs which are translated during arrest and/or during embryogenesis (Kronja et al., 2014a). In order to identify genes that are preferentially translated specifically in arrested oocytes, we performed ribosome profiling and mRNA-seq experiments on 0-2 hour embryos with *Dpse* ovarian extract spike-in and compared these data to those from arrested oocytes. Total normalized ribosome footprints increased 2.7-fold in developing embryos as compared to mature oocytes stored for two days (Fig. 4A). While total translation is lower in arrested oocytes than early embryos, we identified a small subset of “pilot light genes” (243, representing 6.0% of oocyte mRNAs) that are preferentially translated during oocyte arrest (Fig. 4B; Table S1).

**Figure 4.**
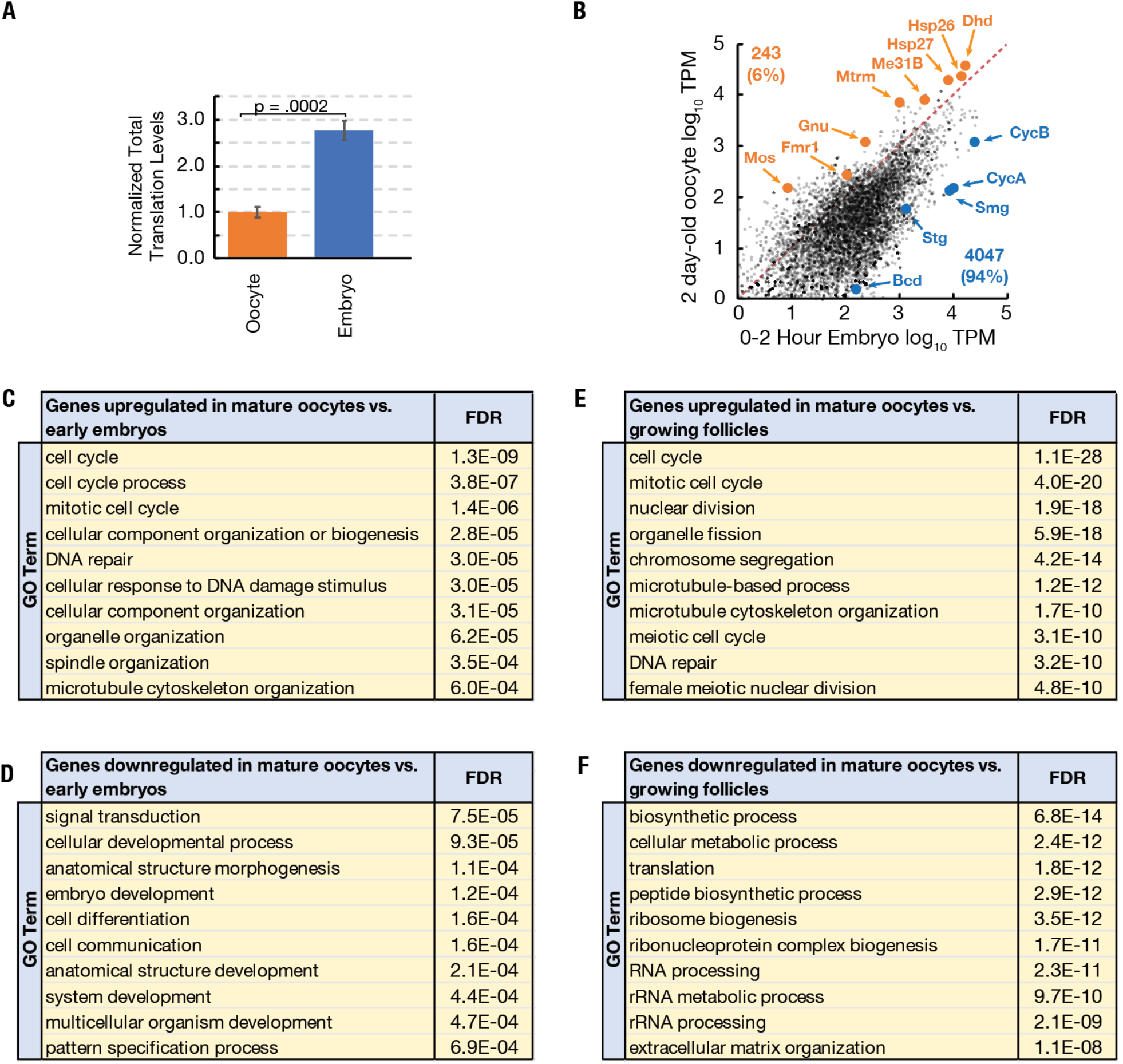
A small fraction of genes are preferentially translated during oocyte arrest. (A) Normalized total translation levels from ribosome profiling is nearly three-fold higher in 0-2 hr embryos compared to oocytes stored for two days. (B) Plot showing most but not all genes are translated at higher levels in the 0-2 hr embryo than the 2 day stored oocyte. Examples of the 243 more highly translated “pilot light genes” are labeled in orange. (C-F) GO analysis of genes with significantly increased (C,E) or decreased (D,F) (p < .01, Student’s t-test) translation in two day old mature oocytes as compared to 0-2 hour embryos (C,D) and growing follicles (E,F).

Many genes preferentially translated in arrested oocytes play important roles in the mature oocyte and shortly after the onset of embryogenesis. For example, *Fmr1* is required to optimally maintain stored oocytes (Greenblatt and Spradling, 2018). Others include the small heat shock proteins *Hsp26* and *Hsp27*, *me31B*, and the thioredoxin-like *dhd*, which is required to remove sperm protamines following fertilization (Emelyanov and Fyodorov, 2016; Tirmarche et al., 2016). In addition, many kinetochore, spindle assembly checkpoint, and meiotic maturation genes including *Ndc80*, *Nek2*, *Zw10*, *gnu*, and *mos* (Lee et al., 2003; Radford et al., 2015; Sagata et al., 1989; Uto and Sagata, 2000; Williams et al., 1996) are also preferentially translated in oocytes (Table S1). GO analysis of the 243 genes showed a significantly enrichment for cell cycle-related processes, including mitotic cell cycle, spindle organization, and DNA repair (Fig. 4C,D).

To investigate whether some functional categories of mRNAs are preferentially stockpiled in advance of oocyte completion, we also gathered information on how gene expression changes as oocyte growth ceases in preparation for storage, ovulation and embryonic development. We carried out RNA-seq and ribosome profiling on the ovaries of young flies 12-16 hours post-eclosion that still lack mature oogenic stages and compared them to day 2 mature oocytes. In addition, we reanalyzed microarray data from hand isolated follicle stages between stage 9 and stage 14 (Tootle et al., 2011) to identify germline expressed genes that are upregulated at stage 10B which should include gene products added to the oocyte just before nurse cell breakdown for storage in the mature oocyte (Table S2).

These studies were consistent with previous analyses of gene expression during late oogenesis and oocyte maturation and showing large increases to genes involved in cell cycle control (Cui et al., 2013; Kronja et al., 2014b; Sieber and Spradling, 2015; Tootle et al., 2011). RNA level changes analyzed by gene ontology reflect completion of follicle growth, nurse cell dumping, reactivation of oocyte meiotic progression from diplotene to a new arrest in metaphase I, and reduced ribosomal production (Fig. 4E, F; Table S2).

### Maintaining meiotic spindles limits oocyte longevity

Given the large reductions we observed in the translation of genes related to meiotic spindle organization and the spindle assembly checkpoint, we investigated whether defects in the meiotic spindle could explain the reduction of oocyte viability during extended storage. We examined 1, 7, and 13 day old *Drosophila* oocytes expressing α-tubulin-GFP, a construct which has been previously used to analyze meiotic spindles (Colombié et al., 2008). We found that the meiotic spindles of day 1 and day 7 oocytes were usually bipolar and highly tapered as previously described (Theurkauf and Hawley, 1992) (Fig 5A, 5A’). By day 13 however, many of the spindles were abnormal, such as unipolar, tripolar or fragmented (Fig 5B-D). Similar defects are seen in mutants of many of the meiotic spindle maintenance genes (e.g. *sub, polo*, and *14-3-3ɛ)* whose translation declined substantially (>2-fold) during oocyte aging. Analyzing embryos derived from aged oocytes, we observed errors in chromosomal segregation during the mitotic divisions that follow pronuclear fusion (Fig. 5F,F’), and meiotic divisions (Fig. 5G,H). In contrast embryos derived from non-aged oocytes progressed normally through cleavage divisions (Fig 5E).

**Figure 5.**
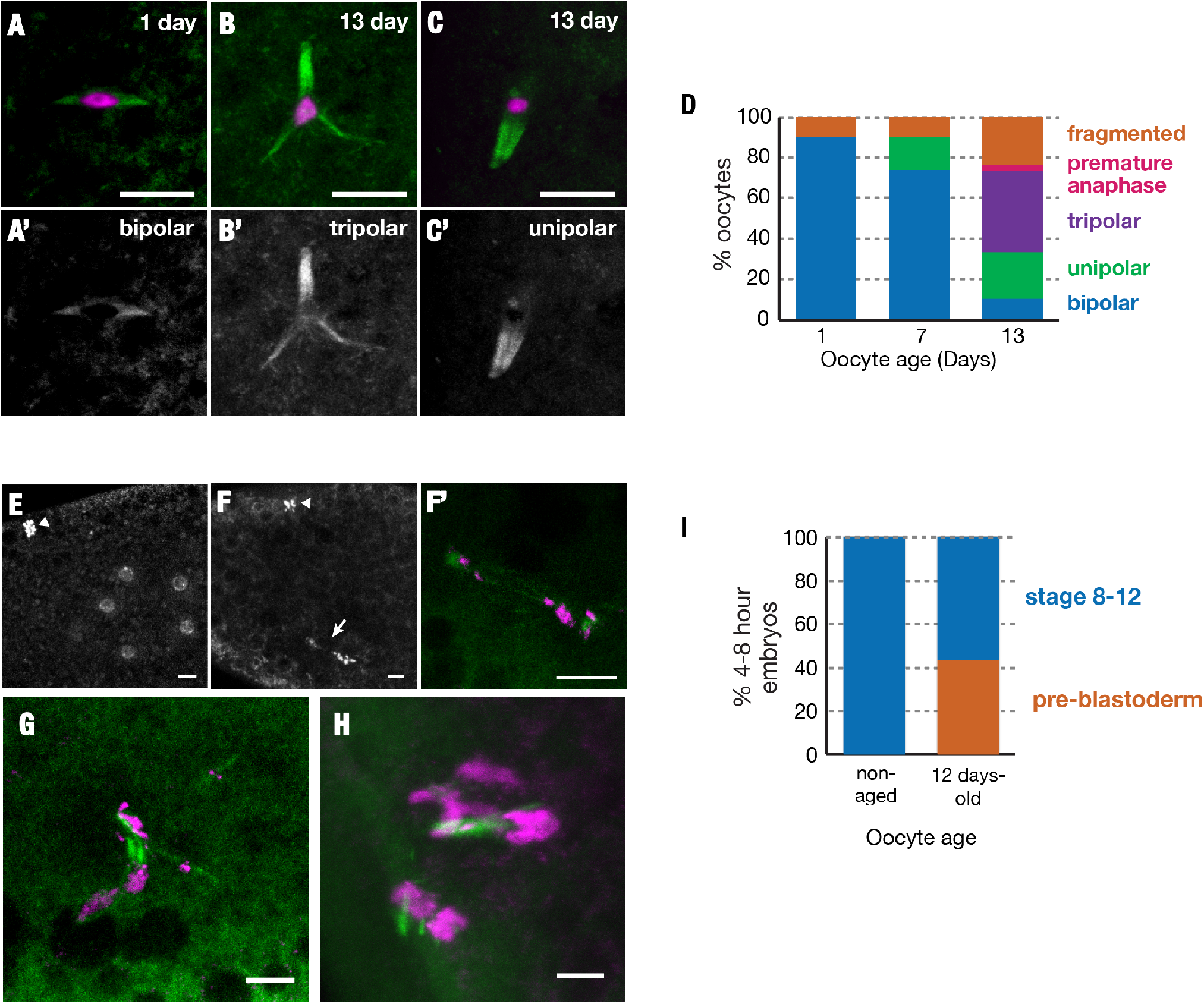
Stored oocytes preferentially fail due to problems of meiotic completion. Meiotic spindles of oocytes stored for 1, 7 or 13 days at 25°C were visualized using α-tubulin-GFP (green) and DAPI (magenta) (A-C), or for α-tubulin-GFP alone (A’-C’). Normal bipolar spindles predominated at 1 day (A), but tripolar (B) and unipolar spindles (C) were readily seen by 13 days. The changes in the frequency of spindle defects is summarized in (D). (E-H) 0-1 hour embryos were collected after adding males to oocytes stored for <1 day or 12 days and analyzed for their developmental state. (E) DAPI stained embryo from a <1 day oocyte with cleavage stage nuclei and condensed polar body (arrowhead) visible at the 8 cell stage, whereas embryos from 12 day old oocytes arrest at the first mitotic (F,F’) or meiotic (G,H) divisions. (F) DAPI stained embryo from a 12 day oocyte is with arrested mitotic spindle (arrow) and polar body (arrowhead). (F’) Zoom-in of the spindle in (F) with tubulin-GFP (green) and DAPI (magenta). (G,H) Chaotic meiosis II divisions in embryos derived from 12 day oocytes with abnormal, tripolar/fragmented spindles. (I) Stage distribution of embryos from non-aged or aged (12 day) oocytes. Embryos fell into two categories; embryos from non-aged oocytes developed to stage 8-12 whereas embryos from 12 day oocytes either progressed to stage 8-12 or arrested during initial meiotic/mitotic divisions (pre-blastoderm). Scale bars = 10 μM.

Only approximately half of embryos derived from oocytes stored for 12 days at 25oC, are able to develop to hatching (Fig. 1D). To investigate whether errors of chromosome segregation are the major cause of reduced embryonic viability, we collected embryos derived from unstored (<1 day) or 12-day old oocytes and analyzed their level of development 4-8 hours after fertilization. 100% of embryos derived from non-aged oocytes had progressed to stages 8-12, as expected for normal development (Fig. 5I). In contrast, the embryos derived from aged oocytes showed a bimodal distribution of development. 60% of these embryos developed to stages 8-12 like young embryos. However, the other 40% arrested during initial cleavage divisions of pre-blastoderm embryos, failing to progress past the mitotic cell cycles of early embryogenesis preceding zygotic genome activation (Fig 5I). This strongly implied that they had undergone chromosome mis-segregation shortly after fertilization leading to lethal aneuploidy. The fact the distribution was bimodal indicates that defects in chromosome segregation arise much more frequently than the many other defects that might cause embryos to arrest at intermediate stages.

### Small heat shock proteins enhance the viability of stored oocytes independently of chromosome segregation

The two small heat shock protein chaperones *Hsp26* and *Hsp27* are among the most highly translated proteins in the mature oocytes, and qualified as potential “pilot light” products since they were translated at higher levels in oocytes than in early embryos (Fig. 4E). Small heat shock proteins (sHSPs) are highly expressed during yeast meiosis (Kurtz et al., 1986), suggesting a potential conserved function for sHSPs during gametogenesis. One potential function of sHSPs might be to stabilize the meiotic spindle and thereby extend the functional lifetime of oocytes. We used FRT-mediated recombination to construct a deficiency strain, *Df(sHSP)*, that eliminates *Hsp26* and *Hsp27* (Fig 6A). The strain was backcrossed 7 times to the *yw* strain to homogenize the genetic background. To determine if any of the other genes in *Df(sHSP)* are expressed in mature *Drosophila* oocyte, we analyzed RNA-seq and ribosome footprint data from wild type and observed that *Hsp26* and *Hsp27* are expressed at high levels, but not any of the other genes (Fig. 6A). We generated antibodies against HSP26 and HSP27, and staining confirmed that they are expressed in developing follicles and strongly expressed at stage 10B shortly before oocyte completion (Fig. 6B).

**Figure 6.**
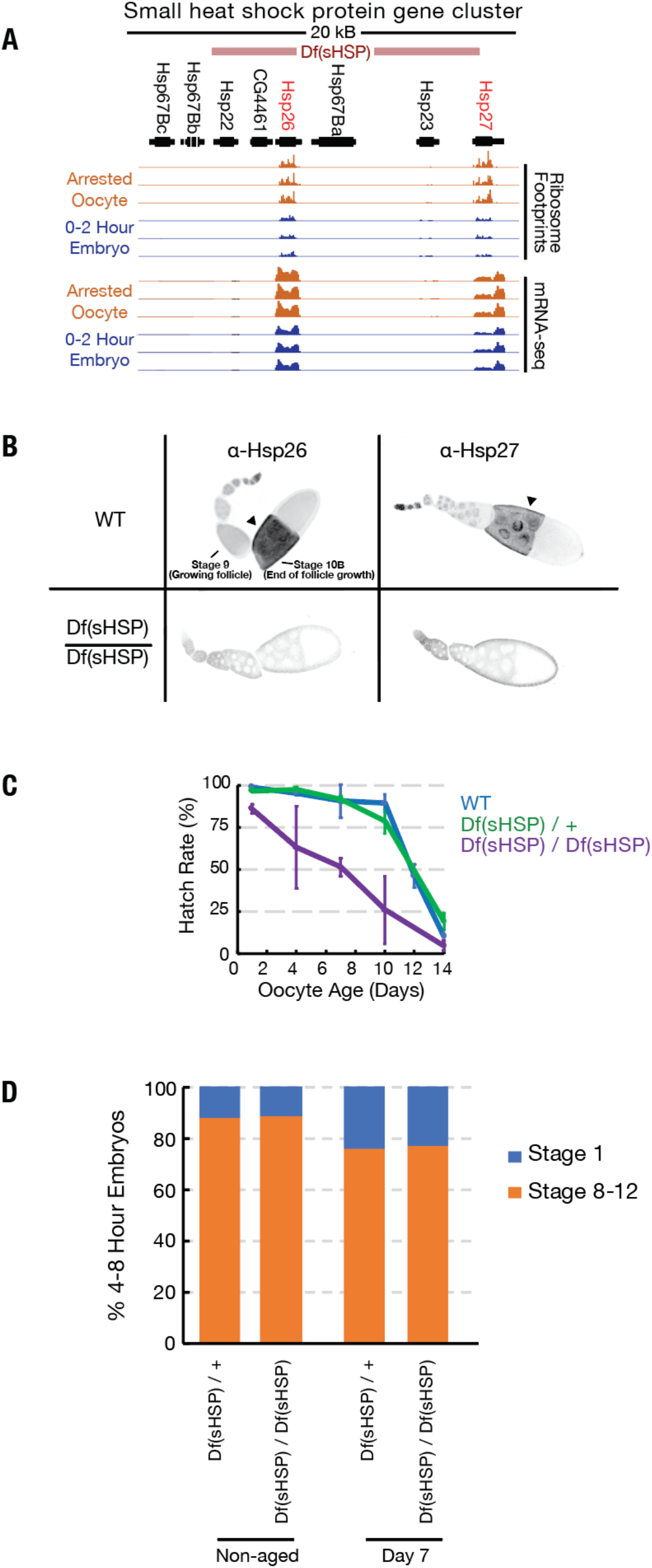
Small heat shock proteins function to maintain quiescent follicles. (A) Normalized ribosome footprinting (upper tracks) and mRNA-seq (lower tracks) read depths in the 67B small heat shock protein gene cluster, are compared for replicate experiments from oocytes stored for 2 day (orange) and for 0-2 hr embryos (blue). Above the tracks is a map of the gene cluster, as well as the extent of a FRT recombination-induced deletion of all but the left-most two genes that was generated. Two genes, *Hsp26* and *Hsp27*, are transcribed and translated in both stored oocytes and 0-2 hr embryos. (B) Wild type (WT) and Df(sHSP)/Df(sHSP) ovarioles were stained with antibodies specific for Hsp26 and Hsp27. Both genes are expressed during oogenesis and are abundant in later stages (stage 10 shown). However, expression was absent above background in stage 10 follicles from the Df(sHSP)/Df(sHSP) ovarioles. (C) The hatch rate of stored wild type (WT) (blue), Df(sHSP)/+ oocytes (green), or oocytes from Df(sHSP) homozygous females (purple) after the days of storage indicated on the x-axis. Deletion of *Hsp26* and *Hsp27* accelerated the rate of decline during storage. (D) Plot showing similar stage distributions of 4-8 hour embryos derived from non-aged or aged Df(sHSP)/+ or Df(sHSP)/Df(sHSP) oocytes, suggesting that the premature loss of viability of oocytes lacking small heat shock proteins is not due to an acceleration of meiotic spindle defects observed in aged wild type oocytes.

The survival of stored wild type oocytes was compared to stored *Df(sHSP)* homozygous oocytes after between 2 and 14 days of storage in vivo to determine if *Hsp26* and *Hsp27* contribute to oocyte stability. Whereas wild type and *Df(sHSP)*/+ oocytes showed normal stability reductions during storage, homozygous Df(sHSP) oocytes lost developmental capacity more quickly (Fig. 6C). However, the reason for the accelerated decline was not due to further destabilization of meiotic chromosome segregation. When the development of embryos derived from non-aged and 7-day-aged *Df(sHSP)* oocytes were examined, the same proportion of embryos arrested early as controls (Fig. 6D). Thus, loss of sHsp chaperones does not strongly affect completion of meiosis and chromosome segregation but must destabilize some later-acting aspect of embryonic development.

## Discussion

### *Drosophila* females can be used to study how mature oocytes age during storage in the ovary

We developed a general system for studying the expression and genetic function of genes involved in the aging of completed *Drosophila* oocytes held in the ovary. Using our approach we determined precise aging curves for mature oocytes and showed they varied with temperature. Identifying the genes required for mature oocyte storage in the absence of transcription will elucidate mechanisms that enhance female fertility in many animals, define the limits of these mechanisms, and provide insight into why rare species such as humans are unable to maintain functional oocytes throughout adulthood.

Our studies also address more fundamental questions about the aging of cells that utilize long-lived mRNAs. Oocytes rely heavily on the regulated translation of relatively stable mRNA populations, especially towards the end of egg production. In this they resemble many other cells, including neurons, that utilize relatively stable mRNA at synapses, and male germ cells, which following meiosis transform into sperm by an elaborate translational program (Besse and Ephrussi, 2008; Fuller, 2016). Normally, an mRNA turns over in a matter of hours, not days (Sharova et al., 2009), and it remains unclear exactly how the functional capacity of mRNAs can be maintained for extended periods. The close association of long-lasting mRNAs in oocytes, neurons and sperm with P bodies, themselves derived from RNA turnover machinery, and the ability of mRNA to cycle between active and inactive states is likely to play critical roles that can now be studied more easily in a relatively simply and tractable *in vivo* system, the mature *Drosophila* oocyte.

### A general genomic analysis of translational changes during aging

Our genomic studies reveal the changes in both mRNA and translation levels of essential *Drosophila* genes throughout the aging process. Our data show that despite the potential instability of mRNAs, a decline in mRNA levels is not involved in the loss of oocyte biological function during aging. Using spike-in controls, it is possible to quantitatively compare samples between different time points. We found no significant decline in mRNA levels over the first 12 days of aging at 25°C.

Despite the preservation of mRNA, there was a pervasive general decline in mRNA translation that correlated with the loss of oocyte function. Translation must decline either because of changes to mRNAs that reduce their ability to be translated, changes to the ribosomes, or alterations in trans-acting factors required for translation. Analyzing the changes in translated proteins during aging did not reveal which of these mechanisms was likely to be responsible. Multiple potentially relevant cellular processes undergo significant decreases in translation. These include genes involved in M phase, in meiotic cohesion, spindle formation, maintenance and bipolarity. Some of these genes are dose sensitive, suggesting that a decrease of two-fold in levels would be enough to generate a phenotype. We observe greatly increased levels of spindle instability as the levels of protein translation fall non-specifically into this range of decline.

### *Drosophila* oocyte decline represents aging in the absence of transcription

An important difference between storage of oocytes within primordial follicles and as full grown oocytes concerns the status of transcription. Primary oocytes can continue to transcribe genes and repair or replace cellular components as needed, while granulosa cells can divide and replace whole cells if necessary. In contrast, mature oocytes without transcription must rely on translation, which despite the presence of sophisticated RNP-based regulatory machinery undergoes a significant decline in translational efficiency over relatively short periods.

What causes the decline in translation over the course of oocyte aging? One possibility is wear and tear on mRNAs that gradually reduces their ability to undergo translation. Even one cleavage usually inactivates an mRNA and targets it for turnover. Most mRNAs decay within a day or less (Sharova et al., 2009) even in growing cells, suggesting that specialized stabilization mechanisms exist during oocyte storage. Unlike proteins, which can be turned over and replaced using mRNA as a template, there is no transcription in mature oocytes and no way to replace damaged mRNA molecules. RNA molecules with expanded trinucleotide repeat sequences can seed the formation of aggregates in a manner analogous to protein aggregation (Jain and Vale, 2011; Querido and Chartrand 2011). It is unknown if mRNAs are generally susceptible to misfolding and aggregation as has been well-characterized for proteins. We hypothesize that P bodies and stress granules, which form during periods of cellular stress as a response to the accumulation of untranslated mRNAs (Eulalio and Izzaurralde et al, 2007), may participate in the long-term preservation of stored mRNAs by preventing mRNA aggregation or oxidative damage.

### Our findings suggest new insight into the strategy of oocyte maintenance in mammals

Our studies have several possible implications for understanding and potentially mitigating the increasing instability of chromosome segregation in mammalian and especially in human oocytes and early embryos. Because of the decades long delay between the onset of meiosis and its completion, some slow decay of an important meiotic process during the primordial follicle stage has been suspected. A logical candidate is the process of sister chromatid cohesion After forming during pre-meiotic S phase, meiotic cohesion complexes do not appear to turn over or incorporate freshly synthesized protein subunits (Revenkova et al., 2010; Tachibana-Konwalski et al., 2010). Reduced levels of the meiotic cohesin complex components Rec8 and Sgo2, which protect cohesin from separase-mediated cleavage, were observed in aged oocytes (Chiang et al., 2010; Lister et al., 2010) and interkinetochore distances increase with age (Merriman et al., 2013).

However, our results suggest that defects arising during the storage of fully grown oocytes have been under-appreciated as an additional source of meiotic and early embryonic mitotic instability. The production of new transcripts in the oocyte GV strongly drops or ceases after oocytes reach full size, and does not begin again until the four cell stage in humans. During this period, oocytes would be largely or entirely dependent on their existing mRNA pool, like stored *Drosophila* mature oocytes. Currently, there are insufficient studies using cell marking techniques to follow how long individual full-size mammalian oocytes remain in a quiescent state, what the consequences of late storage are on translation, and whether the average length of mature oocyte storage changes with maternal age and increased incidence of menstrual irregularities. Given the high sensitivity of mature *Drosophila* oocytes to storage shown here, we suggest that a significant fraction of human chromosome instability is caused by the duration of late storage, rather than by defects that occur at the primordial follicle stage. This would imply that the problems of human chromosome instability may be more susceptible to intervention than previously believed.

## Experimental Procedures

### Oocyte aging assay

Newly eclosed virgin wild type females of indicated genotypes were placed in standard food vials containing added yeast paste made by mixing live yeast with water until the mixture acquires the consistency of peanut butter, but does not trap flies. After feeding on the yeast for 24 hours flies were transferred to plates containing agar-sugar medium to provide humidity, but no edible yeast. After 1 additional day, ovarioles contain two stage 14 eggs, and are considered to have begun day 1 of quiescence. For study, ovaries were dissected after the desired period of quiescence and the mature stage 14 oocytes were collected. To study oocyte viability, 10 females were transferred to chambers with molasses egg laying plates and 10 males were added. Molasses plates were scored with a needle to increase the number of eggs laid. Males were isolated from females for at least 2 days prior to addition and were aged for 3-8 days from eclosion. Laid embryos were counted and recovered for study after various periods of time. Eggs were collected 16 hours after addition of males and hatch rates were determined 48 hours after collection.

### Generation of antibodies and deletion alleles of *Hsp26* and *Hsp27*

Antibodies were generated against the C-termini of *Drosophila melanogaster* HSP26 and HSP27 using peptides KLHcarrier-cys-KANESEVKGKENGAPNGKDK and KLHcarrier-cys-APEAGDGKAENGSGEKMETSK respectively (Proteintech) and were used at a concentration of 1:4,000. A deletion of the sHSP region was generated via FLP-mediated recombination of FRT-bearing lines d00797 and d05052 from the Harvard Exelixis collection (Thibault et al., 2004). Deletion of *Hsp26* and *Hsp27* was confirmed by PCR analysis and immunostaining.

### *Drosophila* ovary and embryo immunostaining

Ovaries were hand-dissected in Grace’s Insect Medium (Life Technologies) from flies fed for 3 days with wet yeast paste. Ovaries were fixed in 4% formaldehyde (37% formaldehyde diluted in PBST (0.2% BSA, 0.1% Triton X-100 in 1X PBS) for 12 minutes. Ovaries were incubated with primary antibodies diluted in PBST with gentle agitation overnight at 4C. Ovaries were then washed 3 times in PBST for at least 20 minutes and incubated with secondary antibodies overnight. Ovaries were then washed 3 times with PBST for at least 20 minutes each, and DAPI (1:20,000-fold dilution of a 5mg/mL stock) was added to the last wash.

Embryos were dechorionated for 2 minutes in bleach (50% diluted fresh Clorox bleach). Embryos were fixed for 25 minutes in a 1:1 mixure of fixative (50mM EDTA, 9.25% formaldehyde, 1XPBS buffer) and heptane with gentle agitation. The lower fixative layer was removed and an equal volume of methanol was added. Embryos were devitellinized by shaking vigorously by hand for 4 minutes, removing the heptane layer, and shaking for an addition 1-2 minutes. Embryos were washed three times in methanol and rehydrated in 50% methanol in PBST, and washed three times in PBST. Embryos were blocked for one hour in PBST and then processed as described for ovaries.

### Ribosome profiling and mRNA-seq library preparation

Ribosome profiling and mRNA sequencing was carried out as described in (Greenblatt and Spradling, 2018) with the following modifications. Aged oocytes were defolliculated by treating ovaries with 5 mg/mL collagenase (Sigma-Aldrich) in PBST for 10 min at room temperature with gentle agitation and then washing oocytes three times in PBST. Oocytes were isolated and separated from debris by filtration. Following extraction of defolliculated oocytes or 0-2 embryos with lysis buffer (0.5% Triton X-100, 150mM NaCl, 5mM MgCl2, 50mM Tris, pH 7.5, 1mM DTT, 20ug/mL emetine (Sigma-Aldrich), 20U/mL SUPERaseIn (Ambion), 50uM GMP-PNP (Sigma-Aldrich)) *Drosophila melanogaster* oocyte extract containing 80ug RNA was combined with *Drosophia pseudoobscura* whole ovary extract containing 1.6ug RNA. The combined extract was then processed as in (Greenblatt and Spradling, 2018).

### Ribosome profiling and mRNA-seq data analysis

Analysis of ribosome profiling and mRNA sequencing data was conducted as in (Greenblatt and Spradling, 2018) with the following modifications. For quantification of bulk mRNA/ribosome footprint levels, adaptor-trimmed reads were mapped to a file containing coding sequences of combined *Dmel* and *Dpse* transcripts using Bowtie v2.3.2 and filtered for only uniquely mapping reads (lines containing string “NH:i:1”) and total reads mapping to either *Dmel* or *Dpse* were counted. Unique *Dmel* reads were then re-mapped to the *Dmel* release 6.02 genome with HISAT2 ver2.1.0. Transcripts per million (TPM) values for coding sequences (ribosome profiling) or exons (mRNA sequencing) were obtained using Stringtie v1.3.5 with the files dmel-CDS-r6.02 gtf or dmel-exons-r6.02.gtf used as a reference annotation for ribosome profiling or mRNA sequencing analysis respectively. Ribosome profiling TPM values from 8 day oocytes, 12 day oocytes, and 0-2 embryo samples were then adjusted by factors of 0.595, and 0.427, and 2.76 respectively to account for bulk changes in translation as determined by the ratios of *Dmel* to *Dpse* reads. Of 26,223 potential 30mer footprints from the top 30 translated *Drosophila* genes in oocytes, we found that 25,324 (97%) of sequences contained at least one polymorphism when comparing sequences from orthologous *Dpse* transcripts.

## Acknowledgements

We are grateful to Kamena Kostova, Steve DeLuca, Chenhui Wang, and members of the Spradling lab for support and comments on the manuscript.

## Competing interests statement

The authors declare that they have no competing financial interests.

Correspondence and requests should be addressed to A.C.S.

## SUPPLEMENTAL FIGURE LEGENDS

**Supplemental Figure 1.**
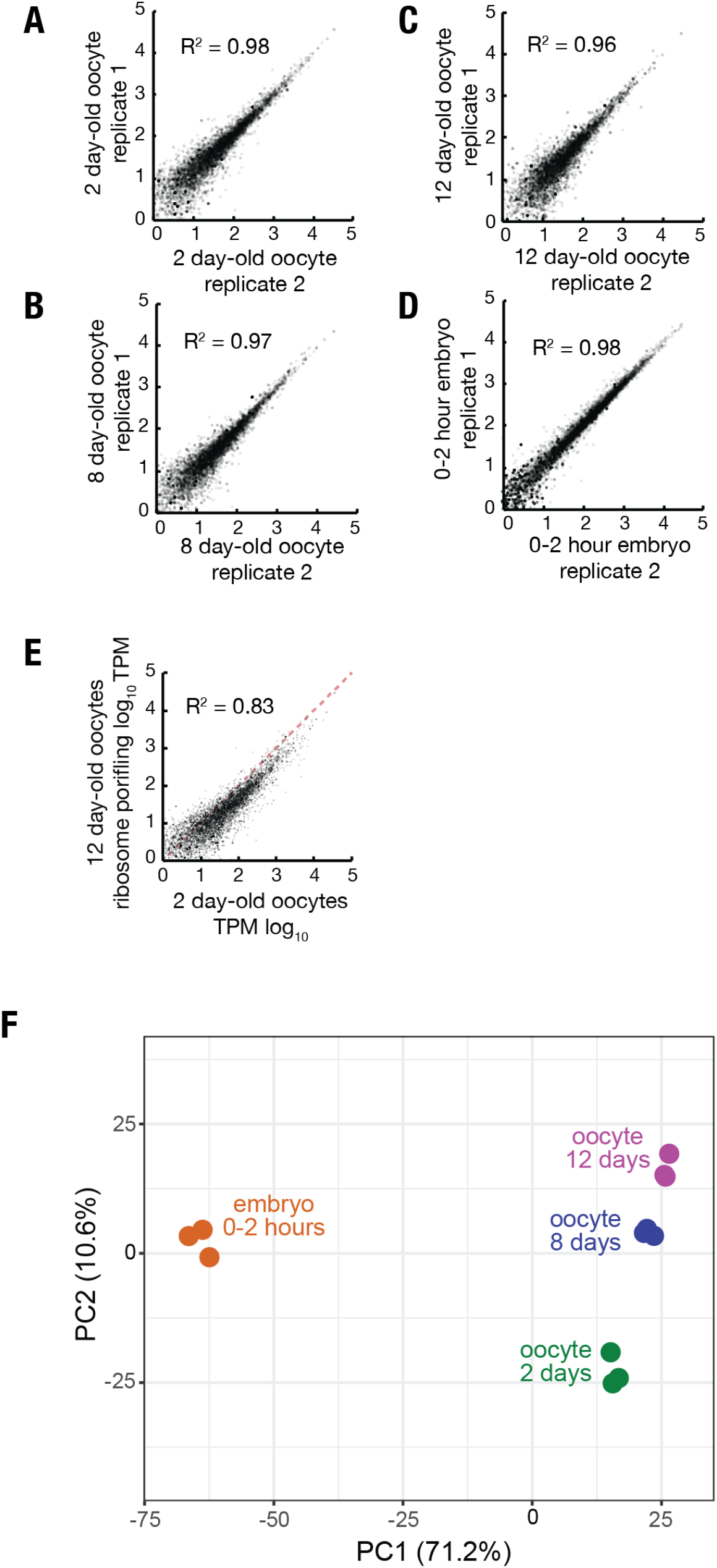
Reproducibility of ribosome profiling data and transcript-length effects on translation efficiency changes in aged mature oocytes. (A-D) Log-log plots showing high reproducibility of individual gene ribosome profiling TPM values from replicate ribosome profiling experiments of 2 day (A), 8 day (B), and 12 day (C) oocytes and 0-2 hour embryos from non-aged oocytes (D). (E) Log-log plot showing that translation of individual genes is less correlated when comparing 12 day to 2 day oocytes versus replicate experiments. (F) Principle component analysis (ClustVis) showing tight clustering of replicate ribosome profiling experiments.

**Supplemental Figure 2.**
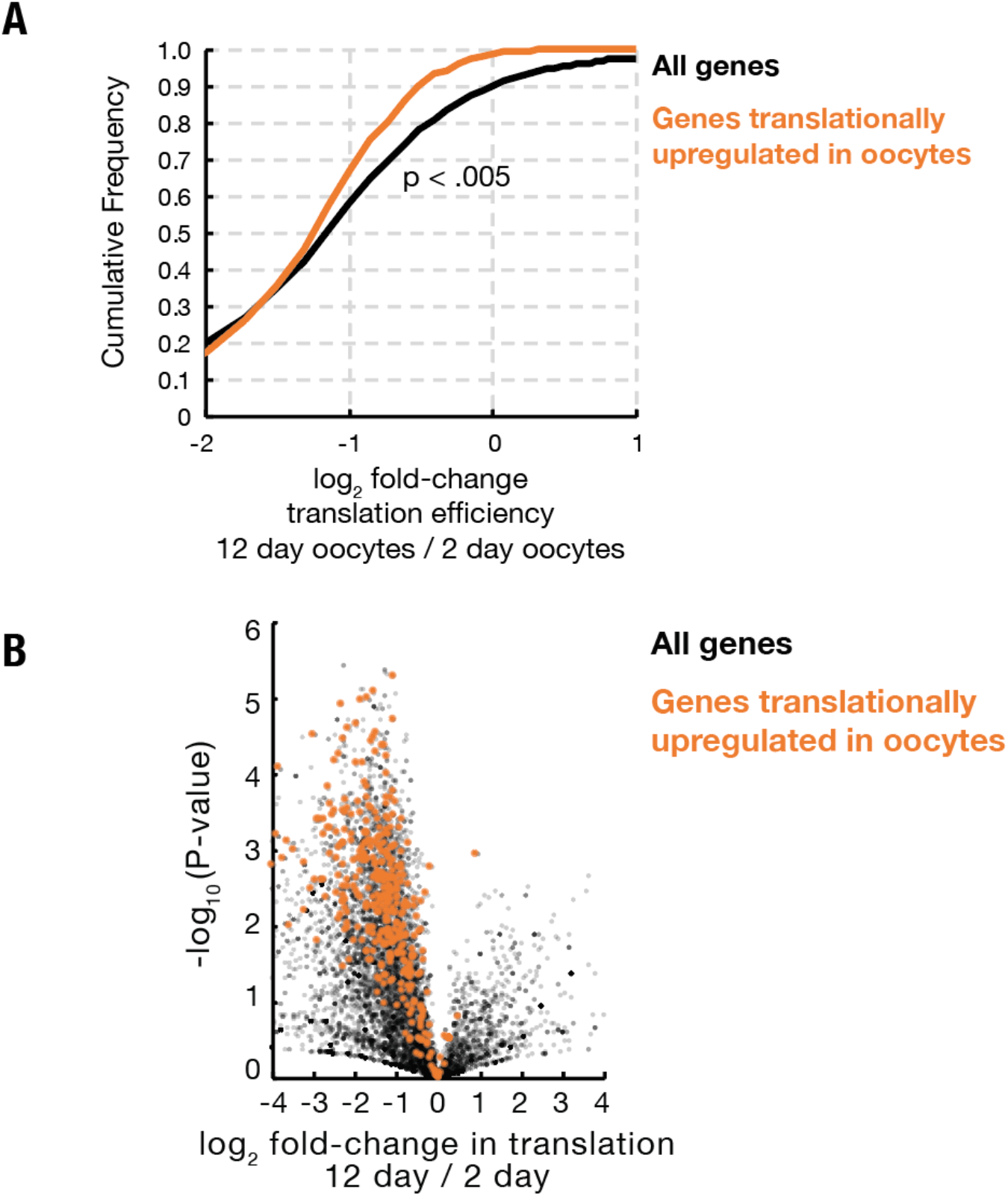
Genes preferentially translated during oocyte arrest are not protected from widespread age-associated reduced translation efficiency. (A) Cumulative distribution plot showing that genes that are translationally upregulated in arrested oocytes (orange) as compared to 0-2 hour embryos show a slightly greater reduction in translation efficiency during aging as a group as compared to the distribution of all genes translated in oocytes. (B) Volcano plot showing that genes preferentially translated in oocytes (orange) as compared to early embryos are globally reduced in translation during oocyte aging.

**Supplemental Table 1. “Pilot light” genes translationally upregulated during oocyte arrest.**

**Supplemental Table 2. Genes upregulated during oocyte maturation.**

**Supplemental Table 3. Counts of total reads mapping to *Dmel* and *Dpse* for ribosome profiling and mRNA-sequencing experiments.**

